# *RiboDiff*: Detecting Changes of mRNA Translation Efficiency from Ribosome Footprints

**DOI:** 10.1101/017111

**Authors:** Yi Zhong, Theofanis Karaletsos, Philipp Drewe, Vipin Sreedharan, David Kuo, Kamini Singh, Hans-Guido Wendel, Gunnar Rätsch

## Abstract

**Motivation:** Deep sequencing based ribosome footprint profiling can provide novel insights into the regulatory mechanisms of protein translation. However, the observed ribosome profile is fundamentally confounded by transcriptional activity. In order to decipher principles of translation regulation, tools that can reliably detect changes in translation efficiency in case-control studies are needed.

**Results:** We present a statistical framework and analysis tool, *RiboDiff*, to detect genes with changes in translation efficiency across experimental treatments. *RiboDiff* uses generalized linear models to estimate the over-dispersion of RNA-Seq and ribosome profiling measurements separately, and performs a statistical test for differential translation efficiency using both mRNA abundance and ribosome occupancy.

**Availability:** *RiboDiff* webpage http://bioweb.me/ribodiff. Source code including scripts for preprocessing the FASTQ data are available at http://github.com/ratschlab/ribodiff.

## 1 Introduction

The recently described ribosome footprinting technology [1] allows the identification of mRNA fragments that were protected by the ribosome. It provides valuable information on ribosome occupancy and, thereby indirectly, on protein synthesis activity. This technology can be leveraged by combining the measurements from RNA-Seq estimates in order to determine a gene’s translation efficiency (TE), which is the ratio of the abundances of translated mRNA and available mRNA [2]. The normalization by mRNA abundance is designed to remove transcriptional activity as a confounder of RF abundance. The TEs in treatment/control experiments can then be compared to identify genes most affected w.r.t. translation efficiency. For instance, [3] considered a ratio (fold-change) of the TEs of treatment and control. However, what these initial approaches only take into account partially is that one typically only obtains uncertain estimates of the mRNA and ribosome abundance. In particular for lowly expressed genes, the error bars for the ratio of two TE values can be large. As in proper RNA-Seq analyses, one should consider the uncertainty in these abundance measurements when testing for differential abundance. For RNA-Seq, this has been described in various ways often based on generalized linear models taking advantage of dispersion information from biological replicates [4, 5, 6]. In [7] and [8], a way to adopt an approach for RNA-Seq analysis for this problem was described that had several conceptual and practical limitations. Here, we describe a novel statistical framework that also uses a generalized linear model to detect effects of a particular treatment on mRNA translation. Additionally, our approach accounts for the fact that two different sequencing protocols with distinct statistical characteristics are used. We compare it to the Z-score based approach [3], *DESeq2* [9] and a recently published tool *Babel* [10] that is based on errors-in-variables regression. Shell and Python scripts for trimming RF adaptor, aligning reads, removing rRNA contamination and counting reads are also included in the *RiboDiff* package.

## 2 Methods

In sequencing-based ribosome footprinting, the RF read count is naturally confounded by mRNA abundance (Fig. 1A). We seek a strategy to compare RF measurements taking mRNA abundance into account in order to accurately discern the translation effect in case-control experiments. We model the vector of RNA-Seq and RF read counts 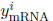 and 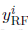, respectively, for gene *i* with Negative Binomial (NB) distributions, as described before [4, 9, 6, for instance,]: *y^i^* ∼ *NB*(*µ^i^*, *κ^i^*), where *µ^i^* is the expected count and *κ^i^* is the estimated dispersion across biological replicates. Here *y^i^* denotes the observed counts normalized by the library size factor (Supplemental Section A). Formulating the problem as a generalized linear model (GLM) with the logarithm as link function, we can express expectations on read counts as a function of latent quantities related to *mRNA abundance β_C_* in the two conditions (*C* = {0,1}), a quantity *β*_RNA_ that relates mRNA abundance to RNA-Seq read counts, a quantity *β*_RF_ that relates mRNA abundance to RF read counts and a quantity *β*_Δ,*C*_ that captures the effect of the treatment on translation. In particular, the expected RNA-Seq read count 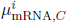 is given by the equation 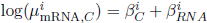.

**Figure 1:**
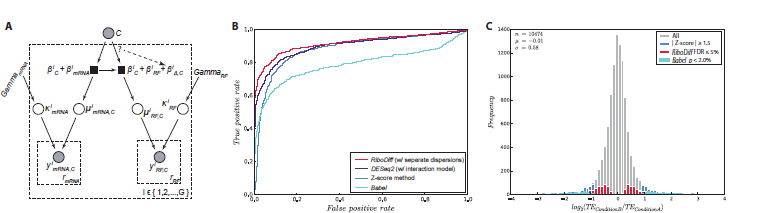
(A) Graphical model representing *RidoDiff* (Gray circle: observable variables; empty circle: unobservable variables; black square: functions; *r* denotes biological replicates; *i* denotes a gene and *G* is the number of genes). The dashed line denotes the relationship that we aim to test (see Methods for details). (B) Receiver operating characteristic (ROC) curve of RiboDiff (with two separate dispersions), *DESeq2* (with interaction model), Z-score method and *Babel* on simulated data with large difference between dispersions of RF and RNA-seq counts (see also Supplementary Figure S-4). (C) Comparison of the distribution of TE ratios of genes that were detected to have a significant change in translation efficiency by *RiboDiff* (w/ joint dispersion), *Z*-score based analysis and *Babel. DESeq2* was very similar to *RiboDiff* (w/ joint dispersion) and was omitted. Data was taken from GEO accession GSE56887.

We assume that transcription and translation are successive cellular processing steps and that abundances are linearly related. The expected RF read count, 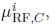 is given by 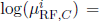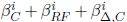. A key point to note is that 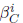 is revealed to be a shared parameter between the expressions governing the expected RNA-Seq and RF counts. It can be considered to be a proxy for shared transcriptional/translation activity under condition *C* in this context. Then, 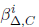 indicates the deviation from that activity under condition *C*, with 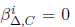 for *C* = 0 and free otherwise (See Supplemental Section B for more details).

Fitting the GLM consists of learning the parameters *β^i^* and dispersions *κ^i^* given mRNA and RF counts for the two conditions *C* = {0,1}. We perform alternating optimization of the parameters *β^i^* given dispersions *κ^i^* and the dispersion parameters *κ^i^* given *β^i^*, similar to the EM algorithm (Supplemental Sections B and C):

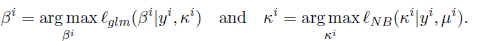

As experimental procedures for measuring mRNA counts and RF counts differ, we enable the estimating of separate dispersion parameters for the data sources of RNA-Seq and RF profiling to account for different characteristics (Supplemental Section E).

As in [5], with raw dispersions estimated from previous steps, we regress all *κ^i^* given the mean counts to obtain a mean-dispersion relationship *f*(*µ*) = *λ*_1_/*µ* + *λ*_0_. We perform empirical Bayes shrinkage [9] to shrink *κ^i^* towards *f*(*µ*) to stabilize estimates (see Supplemental Section D). The proposed model in *RiboDiff* with a joint dispersion estimate is conceptually identical to using the following GLM design matrix protocol+condition+condition: protocol (for instance, in conjunction with *edgeR* or *DESeq1/2*).

In a treatment/control setting, we can then evaluate whether a treatment (*C* = 1) has a significant differential effect on translation efficiency compared to the control (*C* = 0). This is equivalent to determining whether the parameter *β*_Δ,1_ differs significantly from 0 and whether the relationship denoted by the dashed arrow in Fig. 1A is needed or not. We can compute significance levels based on the *χ*^2^ distribution by analyzing log-likelihood ratios of the Null model 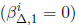 and the alternative model 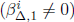.

## 3 Results and Discussion

We simulated data with different dispersions applied to mRNA and RF counts (see Supplemental Section F). We illustrate the performance of our method *RiboDiff* (with separate dispersion estimates) as well as Babel and the Z-score method. Although conceptually closely related to *RiboDiff* with joint dispersion estimates, we also include DESeq2 with a GLM that includes an interaction term (GLM condition + protocol + condition: protocol) to model RNA-seq and RF counts. Figure 1B shows the receiver operating characteristic (ROC) curve for a case with large dispersion differences between RF and RNA-seq counts. *RiboDiff* exhibits a superior detection accuracy compared to *DESeq2*, Babel and Z-score method, which is less pronounced when RF and RNA-seq dispersions are more similar (see Supplementary Figure S-4). We obtained close to identical results for *RiboDiff* with joint dispersion and *DESeq2* with interaction term (data not shown). Our experiments illustrate that it can be beneficial to use the *RiboDiff* model with separate dispersions, in particular, when the dispersions of RF and RNA-seq data differ considerably.

We also re-analyzed previously released ribosome footprint data (GEO accession GSE56887). After multiple testing correction, *RiboDiff* detected 601 TE down-regulated genes and 541 up-regulated ones with FDR ≤ 0.05, which is about twice as many as reported previously. The new significant TE change set includes more than 90% genes identified in the previous study. *RiboDiff* is also compared to Z-score method and we find major differences (see Fig. 1C). Supplemental Section G provides the evidences showing that the Z-score based method is biased towards genes with low read count, whereas *RiboDiff* identifies more plausible differences. *Babel* identifies only very few genes with differential TE. We ran the differential test of *RiboDiff* on a machine with 1.7 GHz CPU and 4GB RAM, it took 23 mins of computing time to finish (10,474 genes having both mRNA and RF counts).

In summary, we propose a novel statistical model to analyze the effect of the treatment on mRNA translation and to identify genes of differential translation efficiency. A major advantage of this method is facilitating comparisons of RF abundance by taking mRNA abundance variability as a confounding factor. Moreover, *RiboDiff* is specifically tailored to produce robust dispersion estimates for different sequencing protocols measuring gene expression and ribosome occupancy that have different statistical properties. The described approach is statistically sound and identifies a similar set of genes from a less developed method that was used in recent work [7]. The release of this tool is expected to enable proper analyses of data from many future RF profiling experiments. The described model assumes that RNA-seq and RF samples are unpaired and it is future work to extend the flexibility of the tool to a broader range of experimental settings.

**Acknowledgements** We thank A. Burcul and & M. Kloft for help. Funding from the Marie Curie ITN framework (Grant # PITN-GA-2012-316861), National Cancer Institute (R01-CA142798-01, H.-G.W.) and the Experimental Therapeutics Center (H.-G.W.).

### A Library Size Normalization

Due to differences in sequencing depth, the read count of the same gene can vary in different samples (or replicates), even if no biological effect exists. Therefore, in a first step, the raw count data needs to be normalized by a library size factor in a first step. We calculate the normalization constant (a.k.a. size factor) S similar to [11] with modifications:

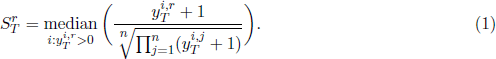

The size factors of RNA-Seq and ribosome footprinting (RF) libraries are calculated separately. Here, *T* denotes data type (RNA-Seq or RF); *r* denotes the r-th sample in data type *T* that includes replicates of both experimental treatments. 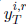 is the observed count of type *T* for gene *i* in sample *r*. For all genes in all replicates, we add one to their count value to avoid the degenerate case of setting the geometric mean across all replicates (indexed by *j*) in the denominator to zero. We calculate the ratios of observed counts of all genes in a given sample to the geometric means and determine the median of these ratios whose count is greater than zero as the size factor. The raw counts of gene *i* in sample *r* are normalized by the corresponding size factor.

### B The Explanatory Matrix of GLM

As described in the main text, we decompose the expected count into multiple latent quantities. For RNA-Seq, it is given by 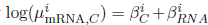, whereas for RF it is given by 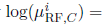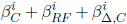. We use a generalized linear model (GLM) to learn the latent quantities *β*s from the observed count data, and then calculate the means. To control the observed read counts fitting into the GLM system, an *n* × 5 explanatory matrix *X* is designed, where *n* equals to the total number of replicates in both experimental conditions for RNA-Seq and RF. Here we show it in the context of linear predictor η of GLM (Equation 2), where *η* = *X* × *β*.

In *X* matrix, the first two columns represent the baseline mRNA abundance 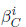 in the two conditions. The third and fourth columns 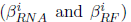 define whether the counts are from RNA-Seq or RF, respectively.^‡^ The fifth column 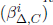 relates the RF count to the potential translational effect. Each row of *X* is used to control how the observed count of a specific sample should be decomposed into latent quantities in order to fit the GLM models. In this example, as indicated by the third column, the first four rows (marked in blue) model RNA-Seq counts with *two replicates* for each condition, *C*0 and *C*1, while the last six rows (marked in green) model RF counts with *three replicates* for each condition. Note the first and second columns in *X* are shared between mRNA and RF counts, where we couple the two different data sets. The linear predictor *η* then is linked with negative binomial distributed mean 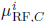 and 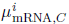 through logarithm as the link function, namely log(*µ*) = *η* = *X* × *β*. The *β*s are estimated by maximizing the likelihood of GLM [12].

In our model, the RNA and RF replicates are assumed to be independent of each other. The flexible configuration allows different numbers of replicates to be analyzed. To extend *RiboDiff* to integrate pairing information, we could modify the GLM model where the pairing term 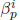(*p* indexes the mRNA and RF pair for each condition) is added to the linear equation. In this way, the pairing signal contributed from a pair of mRNA and RF is absorbed by this term in GLM fitting.

Another possible extension to *RiboDiff* is to handle time series comparison. Assuming we have data at different time points and the goal is to test whether time point *n* is significantly different from others in treatment in a subset of genes. The GLM model for mRNA and RF abundance can be modified as following: 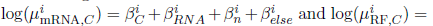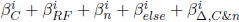, where 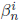 and 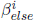 are the latent quantities of time series effects and 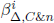 captures the possible TE change at time point *n* under treatment condition.

After the β*^i^* is estimated for each gene, the expected RNA-Seq and RF counts, 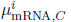 and 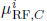 can be obtained. The next step is to estimate the dispersion parameter *κ^i^* by maximizing the negative binomial likelihood function with the observed read counts and the expected counts of both RNA and RF. See details in the next section.

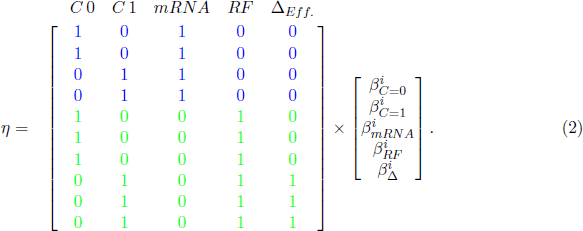

### C Negative Binomial Likelihood Function

Because the count data are assumed to be sampled from a negative binomial distribution with parameter mean *µ* and dispersion *κ*, we estimate *κ* given observed counts and the estimated mean by maximizing the NB likelihood function (equation 4).

The probability mass function of the negative binomial distribution is given by

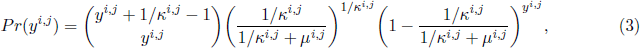

where *y*^*i,j*^ is the observed RF or mRNA read count of *j^th^* replicate of gene *i;* κ^*i,j*^ is the dispersion parameter of the *NB* distribution where *y*^*i,j*^ is drawn from; µ^*i,j*^ is the estimated count of *j^th^* replicate. Thus the logarithmic likelihood of negative binomial of gene *i* is given by

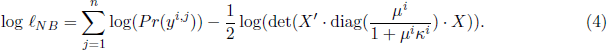

Note that the likelihood function is adjusted by a Cox-Reid term as suggested by Robinson *et al*. [13] to compensate bias from estimating coefficients in fitting GLM step. Again, *X* is the explanatory matrix with dimension *n* × 4 or *n* × 5, depending on *H*_0_ or *H*_1_, where *n* is the total number of RNA-Seq and RF replicates; *µ^i^* is the vector of estimated counts; *κ^i^* is the dispersion vector.

In the previous section, we estimate *β*s by starting with an arbitrary value of dispersion. After we update the dispersion from NB likelihood function, the new dispersion is plugged into the GLM again to start a new optimization cycle. This process ends when the EM-like method converges or an iteration maximum is reached.

### D Empirical Bayes Shrinkage for Obtaining Final Dispersion

From previous steps, we obtain the dispersion for each gene. However, it is estimated only based on the read counts of the gene itself, thus it is less reliable due to the limited replicates. Therefore, we need a systematic method to adjust the raw dispersions. This can be accomplished by the following two steps:

1. Obtain the mean-dispersion relationship by regressing all raw dispersions *κ*^*i*^ given mean counts under assumption: *κ_F_* = *f*(*µ*) = *λ*_1_/*µ* + *λ*_0_ [5]. Namely, for each gene with mean count *µ^i^*, a fitted dispersion 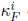 can be calculated.
2. To get the final dispersion 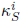, we follow the approach published recently [9]. This approach is based on the observation that the dispersion follows a log-normal prior distribution [14] centered at the fitted dispersion 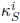. The 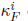 can be estimated by maximizing the following equation:

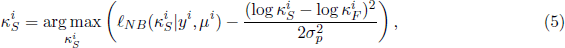

where 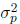 is the variance of the logarithmic residual between prior and the fitted dispersion 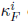. Moreover, the variance 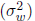 of the logarithmic residual between raw dispersion 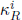 and 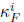 is comprised of 1) the variance of sampling distribution of the logarithmic dispersion 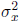 and 2) 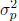. The 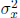 can be approximately obtained from a trigamma function:

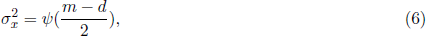

where *m* is the number of samples and *d* is the number of coefficients. Whereas, the 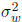 is calculated as the median absolute deviation (mad) of logarithmic residuals between pairs of 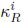 and 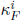:

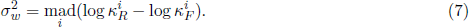

Therefore, we can get the 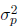 by

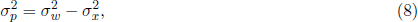

and obtain the final dispersion 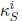 by maximizing the posterior in equation 5.

### E Estimating Dispersion for Different Sequencing Protocols Separately

Because RNA-Seq and ribosome footprinting are different sequencing protocols, the properties of the read counts from these two protocols can vary. Therefore, we enable *RiboDiff to* infer dispersion parameters *κ* for different data sources. Here we show an example where estimating *κ* separately may be needed. The example data are from a recent publication [15].

The empirical dispersion estimates for RNA-Seq and RF counts are calculated from the following equation [9, 4, 11, 16]:

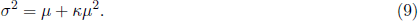

Fig. 2 shows the mean-dispersion relationship. It demonstrates the deviation of the empirical dispersion of RNA-Seq and ribosome footprint data in this experimental setting. The deviation between these two data sets becomes small when read count increases.

**Figure 2:**
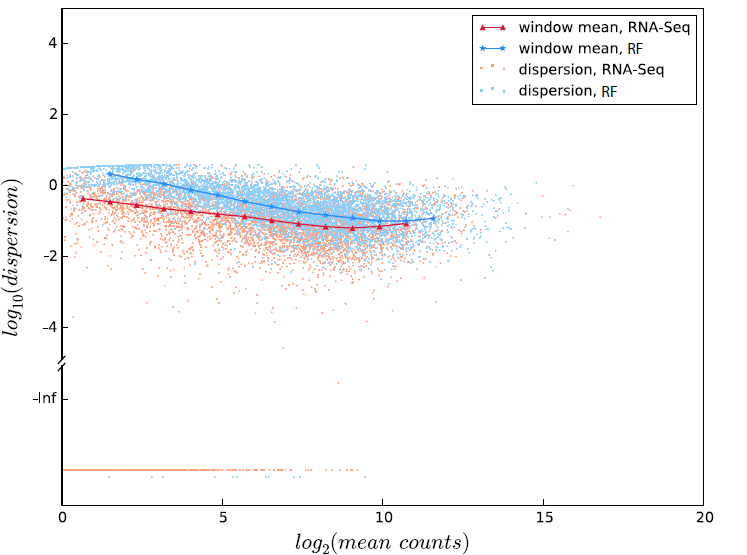
Scatter plot of empirical dispersions. The X-axis is split into several bins and the median *κ* in each bin is highlighted and connected. The empirical *κ* smaller than zero are plotted at the bottom of the figure.

### F Data Simulation

To test the performance of *RiboDiff* and compare it to other methods, we simulated the RF and RNA-Seq read count for 2,000 genes with 500 genes showing down regulated translation efficiency (TE) and 500 genes showing up regulated translation efficiency. There are three replicates for each of the two conditions (i.e., treatment and control) for RNA-Seq and RF. Therefore, count matrix dimensions are 2,000 × 12.

We first generated the mean counts for two treatments of both RF and RNA-Seq across all 2,000 genes assuming their mean counts are randomly drawn from a negative binomial distribution with parameter *n* and *p*, where *n* = 1/*κ* and *p* = *n*/(*n* + *µ*). Then, for each mean count *µ^i^*, we generated three count values as three replicates from a negative binomial distribution with parameter *µ^i^* and *κ^i^*, where *κ^i^* is calculated as *κ^i^* = *f*(*µ^i^*) = *λ*_1_/*µ^i^* + *λ*_0_. To simulate the genes with TE changes in two treatments, we multiply the fold difference to the mean count of the target genes, assuming the fold changes follow a gamma distribution that is observed from real data (GEO accession GSE56887). The gamma distribution has a shape parameter *α* and a scale parameter *s*, and its mean *µ_G_* = *α* · *s*. In the following simulation, we fix *s* and only change *α* to obtain different means for the two treatments and simulate genes having different fold changes using these two means. The fold increase *F_I_* is obtained by

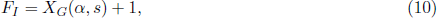

where *X_G_* is a random vector containing 500 elements generated from a gamma density function. And the fold decrease *F_D_* is obtained by

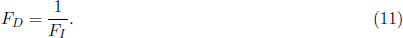

Here, we simulated five groups of count data. In each group, 1,000 out of 2,000 genes showing TE changes:

- mean count has a fold change only for RF count, with *α* = 0.8;
- mean count has a fold change only for mRNA count, with *α* = 0.6;
- mean count has a fold change only for RF count, with *α* = 1.5;
- mean count has a fold change only for mRNA count, with *α* = 1.5;
- mean count has a fold change for RF with *α* = 0.8 AND for mRNA with *α* = 0.6, referred as “combined” in Fig. 3.

**Figure 3:**
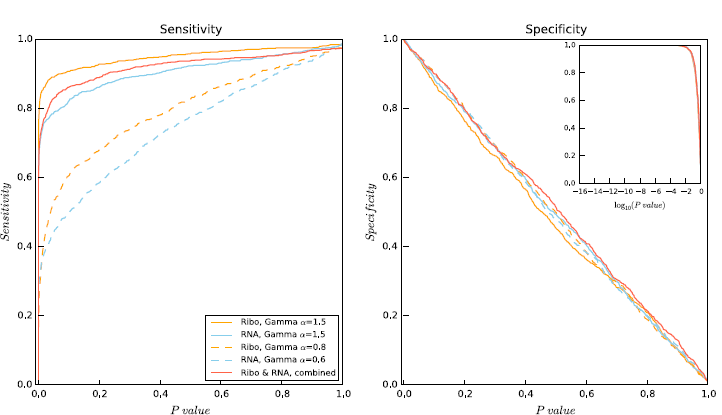
Sensitivity and specificity of RiboDiff on simulated data.

Note that in the last group, if the gene has fold increase in RF, it must have a fold decrease in RNA-Seq. By doing this, the effect at the mRNA level is added to the TE change outcome instead of offsetting the effect caused by RF. Other simulation parameters are as follow: for all RF and RNA-Seq, *n* = 1, *λ*_1_ = 0.1, *λ*_0_ = 0.0001, *s* = 0.5. The parameter *p* controls the scale of the count. We use 0.008 for RF and 0.0002 for mRNA. We run *RiboDiff* with the five dataset to estimate its sensitivity and specificity (Fig. 3).

To evaluate how the number of replicates influences the dispersion estimation, RF and RNA-Seq counts for 5,000 genes with two to ten replicates for each condition were simulated using the same way as described above. For instance, two replicates for condition A and two replicates for condition B in RF, and the same number of replicates for condition A and B in RNA-Seq. In total, we have 9 data sets, and each of them has a certain number of replicates ranging from two to ten. Next, we run *RiboDiff* on these 9 data sets:

*RiboDiff* firstly estimates the raw dispersion *κ^i^* for each gene based on their RF and RNA-Seq counts. Then, a mean-dispersion relationship *κ_F_* = *f*(*µ*) = *λ*_1_/*µ* + *λ*_0_ is obtained by regressing the raw dispersion *κ^i^* given the mean count *µ^i^* using GLM to learn *λ*_1_ and *λ*_0_. Fig. 4A shows the mean-dispersion relationship function for different number of replicates. From this plot we can see that the estimated mean-dispersion relationships, using three to ten replicates, are rather similar to each other, whereas the result using only two replicates deviates from the rest. This indicates that the raw dispersion *κ^i^* estimated using two replicates is less reliable. We observed that the dispersion estimates of high read count genes are larger if only two replicates are used, which can decreases true positive rate.

**Figure 4:**
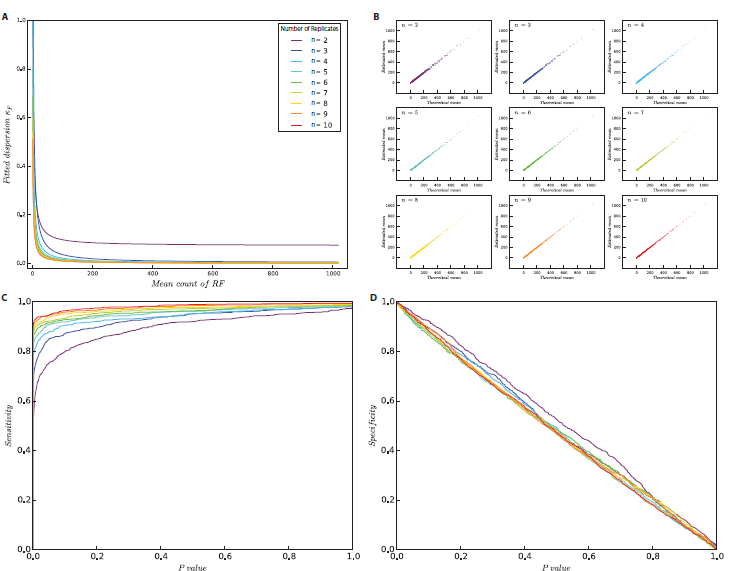
Evaluation of *RiboDiff* by using different number of replicates. (A) Mean-dispersion relationship. (B) Comparison between the theoretical mean and estimated mean. (C) and (D) Sensitivity and specificity of *RiboDiff* calculated under different number of replicates.

We use the same simulated data set to show how the number of replicates affects the latent quantity *β*. For each gene, there are multiple *β*’s that represent different latent quantities, and these *β*s are summed up to obtain the estimated counts of RNA-Seq or RF. Hence, we compare the estimated RF count 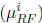 of every gene *i* against their mean counts (theoretical means) that are used to generate the negative binomial counts in the data simulation. In Fig. 4B, each subplot is the comparison of estimated counts (Y axis) from *n* replicates against the theoretical means (X axis). As we can see, the theoretical means and the estimated means correlate well in all 9 experiments (all *r* > 0.99).

Fig. 4C and D show how sensitivity and specificity depend on a chosen *p*-value threshold. For the sensitivity, the area under curves for 2 to 10 replicates increases when the number of replicates increase, whereas the specificities from the same data set do not have large difference among them. This illustrates that the test is well-calibrated and that one can recommend using three replicates to achieve a close-to-best sensitivity.

As *RiboDiff* uses similar technical concept as *DESeq2* [9], we compare the performances of the two methods. Here, *DESeq2* uses a specific design formula: condition + protocol + condition:protocol. The interaction term between sequencing protocol and experimental condition represents the possible condition differences controlling for protocol type.

We simulated three data sets of RNA-Seq and RF counts where gradient differences of dispersion between mRNA and RF were applied to the two data types. The same simulation strategy was used as we described before with modifications. Briefly, 1,000 out of 2,000 genes were chosen to show ΔTE fold change by altering their mean counts of mRNA and RF. The following parameters were used to generate the mRNA count: *n* = 1, *p* = 0.5 × 10^-4^, *λ*_1_ = 0.1, *λ*_0_ = 0.1 × 10^-3^, *α* = 0.8, *s* = 0.5. And for RF count, we used *n = 1, p* = 0.1 × 10^-2^, *λ*_1_ = 10.0, *λ*_0_ = 0.01, *α* = 0.8, *s* = 0.5. Next, we multiplied the mRNA dispersion of every gene in the first data set by a factor of 10, and used the new dispersion to generate mRNA counts for the second data set. Similarly, to obtain the third data set, the original mRNA dispersions were multiplied by 100. We used the same parameters to generate the RF counts for all three data sets. In Fig. 5, from the top to the bottom, the three dispersion plots on the left side show the three simulated data sets where mRNA dispersions are approaching to merge with RF dispersions. The ROC curves on the right side are the corresponding performances of *RiboDiff* with joint and separate dispersion estimates and *DESeq2*. Although *RiboDiff* with joint dispersion estimate performs similar to DESeq2, estimating dispersion separately yields better results under the condition of different dispersions of the two protocols.

**Figure 5:**
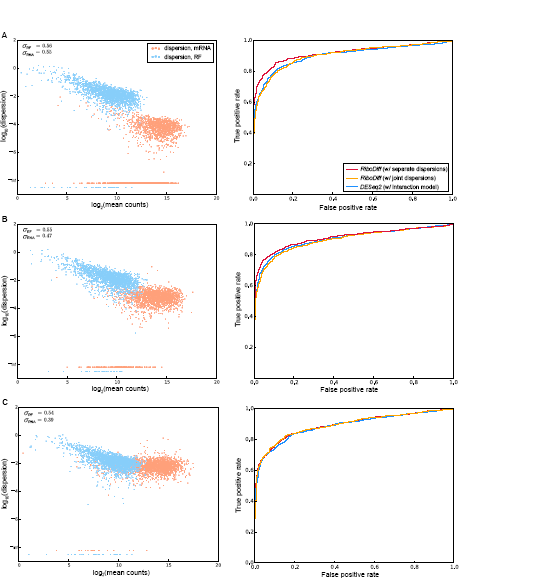
Comparison of ROC curves of *RiboDiff* and *DESeq2* using simulated data. (A-C) The left panel are the dispersions of mRNA and RF; the right panel are the corresponding ROC curves. From the top to the bottom, the differences of dispersion are large, moderate and small, respectively.

### G Results from Real Biological Data

We use previously published ribosome footprint and RNA-Seq data (GEO accession GSE56887) to compare *RiboDiff* with a Z-score based method [3]. The sequencing data were processed in a similar way as before [7], which includes trimming the adapter tail in the reads, aligning the reads, filtering the ribosomal RNA contamination, and counting the reads for genes, *etc*. For gene *i*, the change of translation efficiency Δ*TE^i^* is calculated by

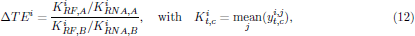

where *t* denotes the data type, as *t* = {RF, RNA-Seq}; *c* denotes the treatment condition A or B, as *c* = {*A*, *B*}; *j* indexes the replicates. 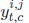 means the *t* type of read count *y* of gene *i* in its *j^th^* replicate under condition *c*. A Z-score was then calculated for each gene as following:

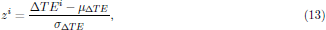

where *µ*_Δ*TE*_ is the mean of Δ*TE* of all genes; σ_Δ*TE*_ is the standard deviation. The genes with |*z^i^*| ≥ 1.5 are selected as significant. Fig. 6A and B show the overlap of significant genes between *RiboDiff* and Z-score based method are limited in both TE down and up regulated gene sets. Further analysis indicates most of the significant genes detected by the Z-score based method having their mean RF counts smaller than 100 with only a few exceptional cases. In contrast, the significant genes detected by *RiboDiff* scatter over a wide range of mean RF count (Fig. 6C and D). It is rational that for highly translated genes, it is more confident to identify significant TE change between two treatments due to enough supported read counts. This is the reason that *RiboDiff* can detect highly translated genes as significant ones even though their absolute value of Z-score are less than 1.5 (|Δ*TE*| below the dashed lines in Fig. 6C and D). This comparison indicates *RiboDiff* identifies more sensible hits and is not biased towards genes with low mean count that inherently have more uncertainty rather than statistically significant differences.

**Figure 6:**
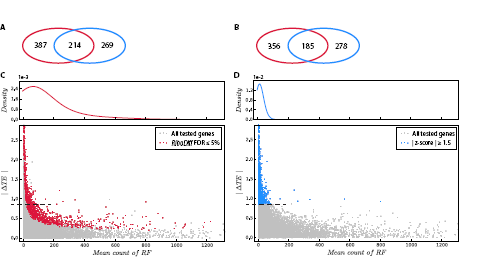
Comparison between *RiboDiff* and a Z-score based method on real biological data. (A and B) Venn diagrams showing the number of overlapping and self specific genes detected by *RiboDiff* and Z-score based method. (A) TE down regulated genes. (B) TE up regulated genes. Red ellipse: results from *RiboDiff*; blue ellipse: results from Z-score based method. (C and D) Scatter plot of mean RF count against the |Δ*TE*|. (C) Result of *RiboDiff*. Significant genes are labeled as red. (D) Result of Z-score based method. Significant genes are labeled as blue. The narrow panels above the scatter plots are the estimated density functions of significant genes on x-axes by using non-parametric kernel density estimation.

Here, we also compare the new TE change gene sets detected by *RiboDiff* and the previous corresponding gene sets published in [7] (Fig. 7). *RiboDiff* detected twice as many as before, and more than 90% genes from the old study are included in the new gene sets.

**Figure 7:**
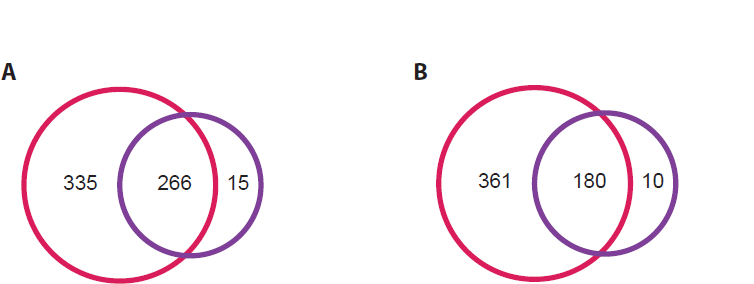
Comparison of results from *RiboDiff* and Wolfe *et al*. (A) TE down regulated genes. (B) TE up regulated genes. Red circle: results from *RiboDiff*; purple circle: results reported in previous study [7].

### H Pipeline cessing of Ribosome Footprinting and RNA-Seq Data Processing

The deep sequencing based ribosome footprinting has many unique features compared to regular RNA-Seq data. Here we discuss its distinct features at each step of data processing before running *RiboDiff’*s differential test on the count input. The relevant scripts that specifically aim for doing each data processing tasks can be found in the *RiboDiff* package.

- Sequencing procedure can introduce bias to the library of replicates. It is always helpful to check the quality of the FASTQ file. Statistics on sequence GC content, length distribution, duplication level, adaptor content and kmer enrichment can provide information from different angles to identify outliers from usable libraries. Publically available tools for doing these tasks are well established and can easily be obtained online.
- In the raw FASTQ file of footprinting, rRNA contamination can take up 25 to 70% of the entire sequences of a library. We construct rRNA databases for specific organisms by collecting their rRNA sequences from SILVA [17]. Both footprint and RNA-Seq reads are aligned to the rRNA database to identify rRNA reads that need to be filtered.
- Next, we use STAR [18] to align both footprint and RNA-Seq reads to the reference genome to get mapping information. STAR is an ultrafast and accurate aligner that also supports junction reads crossing exon splicing site. It also trims the linker sequence (CTGTAGGCAC-CATCAAT) on the 3’ of footprint reads while aligning them to the reference. Ribosome protected mRNA sequences are short (normally from 20 to 40 nt), therefore, to minimize the effect of multiple mapping, only uniquely aligned reads are used.
- The identified rRNA reads from step 2 are removed from the alignment of footprint and RNA-Seq. It is also recommended to check whether footprint reads are clipped even after trimming the linker sequence. Over-clipped short reads are prone to produce ambiguous alignments.
- The last step of preprocessing the data is counting reads for each gene. We add a counting script that only takes the reads mapping the exonic regions guided by an annotation GTF file.

## Footnotes

‡ In the implementation, in order to keep full rank of *X*, we do not include the fourth column 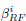, as it is linearly dependent with the third column.

